# Quantification of membrane fluidity in bacteria using TIR-FCS

**DOI:** 10.1101/2023.10.13.562271

**Authors:** Aurélien Barbotin, Cyrille Billaudeau, Erdinc Sezgin, Rut Carballido-López

## Abstract

Cell membrane fluidity is an important phenotypic feature that regulates the diffusion, function and folding of transmembrane and membrane-associated proteins. It is particularly interesting to study it in bacteria as variations in membrane fluidity are known to affect fundamental cellular processes such as respiration, transport and antibiotic resistance. As such key parameter, membrane fluidity is regulated to adapt to environmental variations and stresses like temperature fluctuations or osmotic shocks. Membrane fluidity has been however scarcely studied quantitatively in bacterial cells, mostly because of the lack of available tools. Here, we developed an assay based on total internal reflection fluorescence correlation spectroscopy (TIR-FCS) to directly measure membrane fluidity in live bacteria via the diffusivity of fluorescent membrane markers. We used this assay to quantify the fluidity of the cytoplasmic membrane of the Gram-positive model bacterium *Bacillus subtilis* in response to a cold shock, caused by a shift from 37°C to 20°C. In our experimental conditions, steady-state fluidity was recovered within 30 mins, and the steady-state fluidity at 20°C was about half of that at 37°C. Our minimally invasive assay opens up exciting perspectives and could be used to study a wide range of phenomena affecting the bacterial membrane, from disruption by antibiotics, antimicrobial peptides, or osmotic shocks.

**Significance:** Using fluorescence correlation spectroscopy (FCS) with total internal reflection fluorescence (TIRF) illumination, we measured the diffusion speed of fluorescent membrane markers as a readout for membrane fluidity of growing *B. subtilis* cells. Quantification of the effect of cold shock provided unique information about the dynamics of the plasma membrane of *B. subtilis*. The unprecedented capability of TIR-FCS to quantify membrane fluidity in living bacteria opens the door to a whole set of new studies that will shed light on the bacterial plasma membrane and its interactions with the environment.

## Introduction

The plasma membrane is a component of virtually every living cell, made of a fluid mixture of lipids and proteins, that separates the intracellular and extracellular spaces. The fluidity of the plasma membrane is the physical parameter that defines how fast a given element can diffuse within the membrane at a given temperature. Thus, it is of utmost interest for protein diffusion and biomolecular interactions (1, 2). Moreover, membrane fluidity affects protein folding (3–5). In bacteria, membrane fluidity was found to to be critical in both Gram-negative (for respiration in *Escherichia coli* (6) and multidrug transport in *Methylobacterium extorquens* (7)) and Grampositive (*e*.*g* resistance to antibiotics in *Staphylococcus aureus* (8)) bacteria. Membrane fluidity in bacteria was found to vary in response to chemical (9) biochemical (10, 11) or osmotic (12, 13) stresses. Another hint of the importance of membrane fluidity in bacterial cells is the widespread existence of control systems that maintain it by modifying lipid and protein composition (14, 15). In particular, the fatty acid composition of phospholipids – the main class of lipids of the plasma membrane – is modified in response to changes in temperature to modulate steric constraints and thereby lipid packing (16). Furthermore, proteins such as flotillins (17, 18) and MreB (19) in *B. subtilis* are also thought to play a role in membrane fluidity. In the case of extremophiles, the generation of exopolymers (20), cryoprotectants and antifreeze proteins (21) are used by bacteria to protect their membrane against changes in temperature. Existing assays to measure membrane fluidity in live and synthetic membranes include electron spin resonance (ESR) (22, 23) and nuclear magnetic resonance (NMR) spectroscopy (17), membrane fatty acid analysis (8, 12, 22), fluorescence assays using environment-sensitive probes like diphenylhexatriene (DPH) or ratiometric probes like Laurdan (9) and the measurement of the diffusion speed of a fluorescent tracer using either single particle tracking (SPT) (24), fluorescence recovery after photobleaching (FRAP) (25) or fluorescence correlation spectroscopy (FCS) (26). In microbiology, the most frequently used techniques are fatty acid analysis and DPH anisotropy or ratiometric imaging. Fatty acids analysis informs on the membrane composition but is an indirect readout of fluidity. It gives multidimensional results (relative proportions of different fatty acid with branching and (poly)unsaturation), which can be challenging to directly associate with a change in membrane fluidity, and can in the best case only provide qualitative comparisons of the resulting fluidity. Environment-sensitive fluorescent probes can give useful insights but can only measure relative differences in membrane fluidity. Many probes exist that are sensitive to different parameters of the membrane (27, 28) and their behaviour can be biased by unforeseen interactions (29). FCS has occasionally been used in a few instances in bacterial membranes, to study protein diffusion (30– 32), RNA concentration (33), assembly of protein complex (34) or membrane dynamics in response to antibiotic treatment (26). These studies were all performed using confocal microscopy which axial resolution is not well suited for measurements in bacteria. The axial resolution of confocal microscopes is comparable to the diameter of most studied bacterial cells (500 nm – 1 µm), and this results in problems such as such as having both top and bottom membranes in focus at once or excessive background from out-of-focus membranes. These limitations can be overcome either by using super-resolution (35) or more simply by using total internal reflection fluorescence (TIRF) microscopy. TIRF significantly improves the axial resolution of a microscope by illumination with an evanescent field that usually significantly decays within a range of 100 nm. Total internal reflection fluorescence correlation spectroscopy (TIR-FCS) (36, 37) was previously used to study molecular dynamics in eukaryotic cells (38, 39) and to measure the fluidity of flat synthetic membranes (40, 41). However, TIR-FCS was not previously applied to bacteria despite TIRF being the standard for membrane investigations in such organisms (42). TIR-FCS offers several advantages over confocal FCS, besides the unrivalled axial selectivity of TIR illumination: camera-based TIR-FCS also offers massive parallelisation of measurements, since hundreds of FCS curves can be acquired at once instead of a single one on a confocal microscope. TIR-FCS can also easily generate diffusion maps and therefore retrieve spatial information. Finally, TIR-FCS enables, by resampling intensity fluctuations in space after acquisition, the measurement of diffusion speeds at different spatial scales (spot-variation FCS (43)). Here, we propose to extend the scope of application of TIR-FCS to measure membrane fluidity in live bacterial cells, exemplified by the Gram-positive leading model organism *Bacillus subtilis*. To simplify data analysis and fully use the high-throughput capability of imaging FCS, we developed a new FCS quality metric to automatically discard artefactual curves. Using simulations validated by experiments in synthetic samples, we measured the bias induced by the small size and curvature of bacterial membranes on TIR-FCS measurements. We demonstrated the validity of our assay by studying a well-known system: the response of the *B. subtilis* plasma membrane to a cold shock (44, 45). Diffusion measurements of the membrane markers Nile Red and Di4-ANEPPS confirmed the previous knowledge about cold shock recovery and provided unprecedented insights of bacterial membrane dynamics at different temperatures.

## Methods

### FCS setup

TIR-FCS acquisitions were performed on a Zeiss Elyra PS1 microscope equipped with a 100×/1.46 NA Apochromat oil immersion objective. A typical FCS acquisition consisted of 50000 frames, on a field of view of 128x10 pixels with a pixel size of 160 nm in the object plane and a frame acquisition time of 1.26 ms, the maximum achievable with our camera. Stable focus was ensured using Definite Focus. Detection was performed using an emCCD camera (Andor iXon), using maximum pre-amplification (5×) and electron-multiplying gains (300×) settings. Laser excitation at 561 nm was set, unless specified otherwise, to 5% of the maximum excitation power, corresponding to a power of 460 µW in epifluorescence mode measured in the focal plane of the objective. The excitation area was of approximately 80x80 µm size, leading to an estimated power density of ∼70 nW/µm^2^.

### Data processing and fitting

Pixels from each image stack were numerically binned 2 by 2, unless specified otherwise. The first 1500 frames of every acquisition were discarded as an occasional loss of focus could lead to artefactual intensity fluctuations in these frames. Intensity timetraces at each binned pixel were corrected for bleaching using a double exponential fit (46). Intensity traces at each pixel were correlated using a python implementation of the multipletau algorithm (47). FCS curves were fitted using the standard 2D imaging FCS model (48) except in bacterial cells where the fitting model is described in the ‘technical implementation’ section:

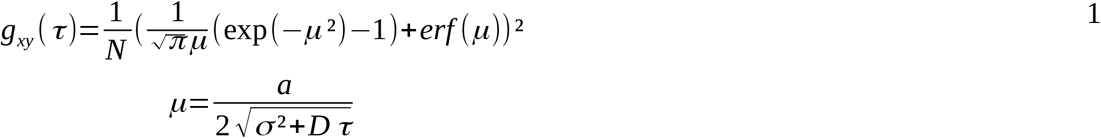

Where *N* is the average number of molecules in the observation area, *a* is the effective pixel size (320 nm with 2x2 pixel binning), *D* the diffusion coefficient, *τ* the lag time and σ is the standard deviation of the microscope’s Point Spread Function (PSF) approximated as a two-dimensional Gaussian function:

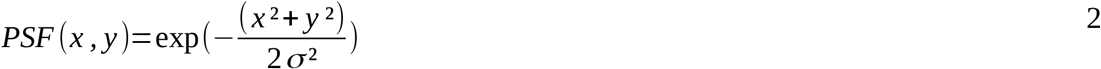

Calibration of the PSF was done as described in reference (49). We measured our PSF σ=0.19 µm, corresponding to a full width at half-maximum (FWHM) of 450 nm, larger than expected using a 1.46 NA oil immersion objective. This enlargement was likely caused by our use of a low magnification tube lens that degraded resolution, as we measured a PSF size σ=0.16 µm when using a higher magnification tube lens. For all acquisitions except in SLBs, we used an intensity threshold set to 80% of the maximum intensity in the field of view (see Supplementary Information for the determination of intensity threshold).

### Liposomes preparation

1,2-di-(9Z-octadecenoyl)-*sn*-glycero-3-phosphocholine (DOPC) and 1-palmitoyl-2-oleoyl-*sn*-glycero-3-phosphocholine (POPC) stored in chloroform were purchased from Merck (Darmstadt, Germany) and stored under argon. 50 µL of 10 mg/mL stock were added to a glass tube then dried using Argon under rotation. Lipids were resuspended in 1.6 mL Phosphate-Buffered Saline (PBS), then tip-sonicated for 10 mins in 30 s on/off cycles on ice. Liposomes were labeled before experiments with 1% of 1,2-dioleoyl-sn-glycero-3-phosphoethanolamine-N-(lissamine rhodamine B sulfonyl) (PE-Rhodamine) (Merck) at a concentration of 10µg/mL.

### Supported lipid bilayer preparation

Supported lipid bilayers (SLB) were prepared by liposome deposition. We pipetted 20µL of liposome solution to a home-made microfluidic chamber made of a sandwich of a slide and a plasma-cleaned coverslip held together by two strips of molten parafilm. Excess liposomes were abundantly washed using 200µL PBS. The chamber was then sealed using parafilm to prevent evaporation.

### Beads-supported lipid bilayer preparation

Beads-supported lipid bilayers were prepared as described elsewhere (50). 10µL of 5 µm uncoated silica beads (BioValley, Nanterre, France) were washed twice in 1 mL PBS, then mixed with 50µL liposomes. The mix was shaken for 20 mins to form BSLBs, then washed twice in 1 mL PBS. 200 µL PBS was left after final wash. 100 µL of BSLBs were then pipetted to a glass-bottom Ibidi chamber for imaging.

### Cells preparation and staining

The wild-type laboratory strain 168 trpC2 of *B. subtilis* was grown and imaged in rich lysogeny broth medium (LB). For measurements at 37°C, 3µL of cells from an overnight culture grown at 30°C were diluted in 2 mL LB and grown at 37°C under agitation for 2h30 until they reached exponential phase (OD_600_ ∼ 0.3) then labeled with 0.2% (v:v) of either 50 µg/mL Nile Red or 40µg/mL Di4-ANEPPS (Thermofisher) disolved in DMSO. Labeled cultures were left under agitation for 15-20 additional min, then 3 µL were transfered to an agarose pad (1.2% in LB) for immobilisation and covered with a plasma-cleaned coverslip.

For steady-state measurements at 20°C, cells from an overnight culture at 30°C were first diluted (3 µL of overnight culture diluted in 2 mL fresh LB) and grown for about 2 hours at 37°C until reaching exponential phase (OD_600_ ∼ 0.2), transferred at 20°C under agitation for 4-5h for at least one generation, then labeled as described above.

When cells were transferred from the liquid culture to the agarose-coated slide, we observed temporary membrane readaptation for about 25 mins (Figure S1). This readaptation could be due to osmotic shock (51), oxidative stress or another cause that remains unknown. We therefore leave cells to settle on the agarose-coated slide for 25 mins on the slide before starting to image. In the case of cold shock, cells were first immobilised on an agarose pad and covered with the coverslip at 37°C for 25 min then transferred at 20°C in the microscope.

## Technical implementation

### Filtering curves based on goodness of fit

FCS measurements can be subject to artefacts that distort FCS curves, for instance when bright clusters of fluorescent molecules enter the observation area. These curves need to be discarded from analysis to avoid biasing diffusion coefficient estimation. In point FCS, this is often done via manual inspection facilitated by dedicated softwares (52). This approach is however impractical in imaging FCS due to the high parallelisation of FCS curves acquisition resulting in the generation of a high number of FCS curves (typically we acquire 350-500 FCS curves per hour in bacteria, considering only one pixel binning value). The issue of sample-induced artefact in FCS is well acknowledged and solutions to this issue were previously developed. It was notably proposed to compare each curve within a dataset to the averaged curve and to exclude outliers (53). This solution is however computationally intensive and might fail if signal levels vary within a dataset, for instance due of cell-to-cell heterogeneity. Another approach consists in rejecting curves with irregular residuals. One method for doing this consists in calculating the χ^2^ goodness of fit (30), but it does not handle well noisy curves (53). It also requires knowledge of the standard deviation of the FCS curve, which is not always available. The Fourier transform was previously used to detect unevenly distributed residuals, relying however on fine-tuning three empirical parameters and making assumptions on the transit times observed (33). As an alternative to these methods, we introduce here a simple error metric based on fitting residuals that quantifies fitting bias. Considering the mean square error (MSE):

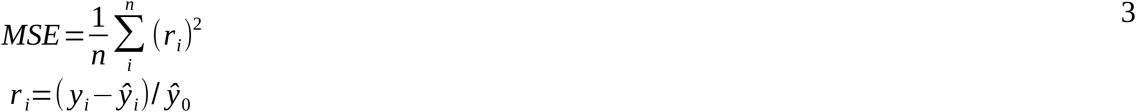

Where *y*_*i*_ is the empirical FCS curve at lag time *I, ŷ*_*i*_ is the corresponding fit value and *r*_*i*_ the residual. A high MSE value is indicative of either a high fitting bias in a poorly fitted curve, which needs to be discarded, or of low signal value (54) caused by strong oscillations of the FCS curve around its fit. Expecting strong variations in signal levels within and across acquisitions, due to cell-to-cell variations and inhomogeneous illumination of curved cells in the TIRF field, we designed a metric that measures fit quality with a lower dependency to signal level. This metric, named non-linear mean-square error (MSE_nl_) is a weighted sum of residuals, where the weight of each residual is equal to the number of adjacent residuals of same sign *n*_*adj,i*_:

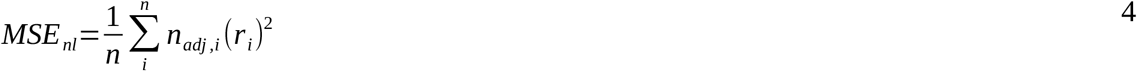

Concretely, if residuals *r*_*i*_ between *i-2* and *i+2* are all positive, the fitting bias is strong and the weight *n*_*adj,i*_=4 is high. On the other hand, if the residual *r*_*i*_ is positive but the residuals *i-1* and *i+1* are negative, there is no fitting bias (residuals are oscillating around the mean) and *n*_*adj,i*_=0. To evaluate the capability of *MSE*_*nl*_ to evaluate fitting bias, we performed 2 TIR-FCS acquisitions on a flat sample of DOPC SLBs, at either high (740 µW) or low (185 µW) excitation power. The resulting dataset had heterogeneous signal levels representing the expected heterogeneity in biological samples. This dataset being acquired on the same SLB however we expected to find a comparable number of artefactual curves with either excitation intensities. Comparing *MSE* and *MSE*_*nl*_ for every FCS curve in the dataset (Figure 1A), we could observe that excitation intensity was a good predictor of *MSE* but not of *MSE*_*nl*_. Curves acquired at higher excitation intensity had in average a lower *MSE* but similar *MSE*_*nl*_, showing a lower dependence of *MSE*_*nl*_ with signal levels. We observed a strong correlation between the two metrics for curves acquired at a same excitation intensity, as poorly fitted curves have higher residuals than well-fitted curves at similar signal levels (Figure 1B). However, comparing two FCS curves with same *MSE* but different *MSE*_*nl*_ suggested that *MSE*_*nl*_ measures fitting bias irrespective of signal levels, unlike *MSE* (Figure 1C-D). Considering that every FCS curve contains imperfections, we then sought to determine the maximum acceptable amount of fitting bias measured by *MSE*_*nl*_. For this, we plotted the measured diffusion coefficient against *MSE*_*nl*_ (Figure 1E, bottom). We observed that high fitting bias were correlated to slower diffusion coefficients, themselves caused by artefacts in FCS curves. We confirmed this by plotting the average diffusion coefficient for sets of FCS curves with *MSE*_*nl*_ thresholds (Figure 1E, top) and found that FCS curves with a low *MSE*_*nl*_ corresponded to the expected diffusion coefficient (55). Using both inspection of individual FCS curves and the scatterplot shown in Figure 1E, we set the *MSE*_*nl*_ threshold to the value of 0.015. We kept this threshold throughout this study, in both synthetic and biological samples.

**Figure 1:**
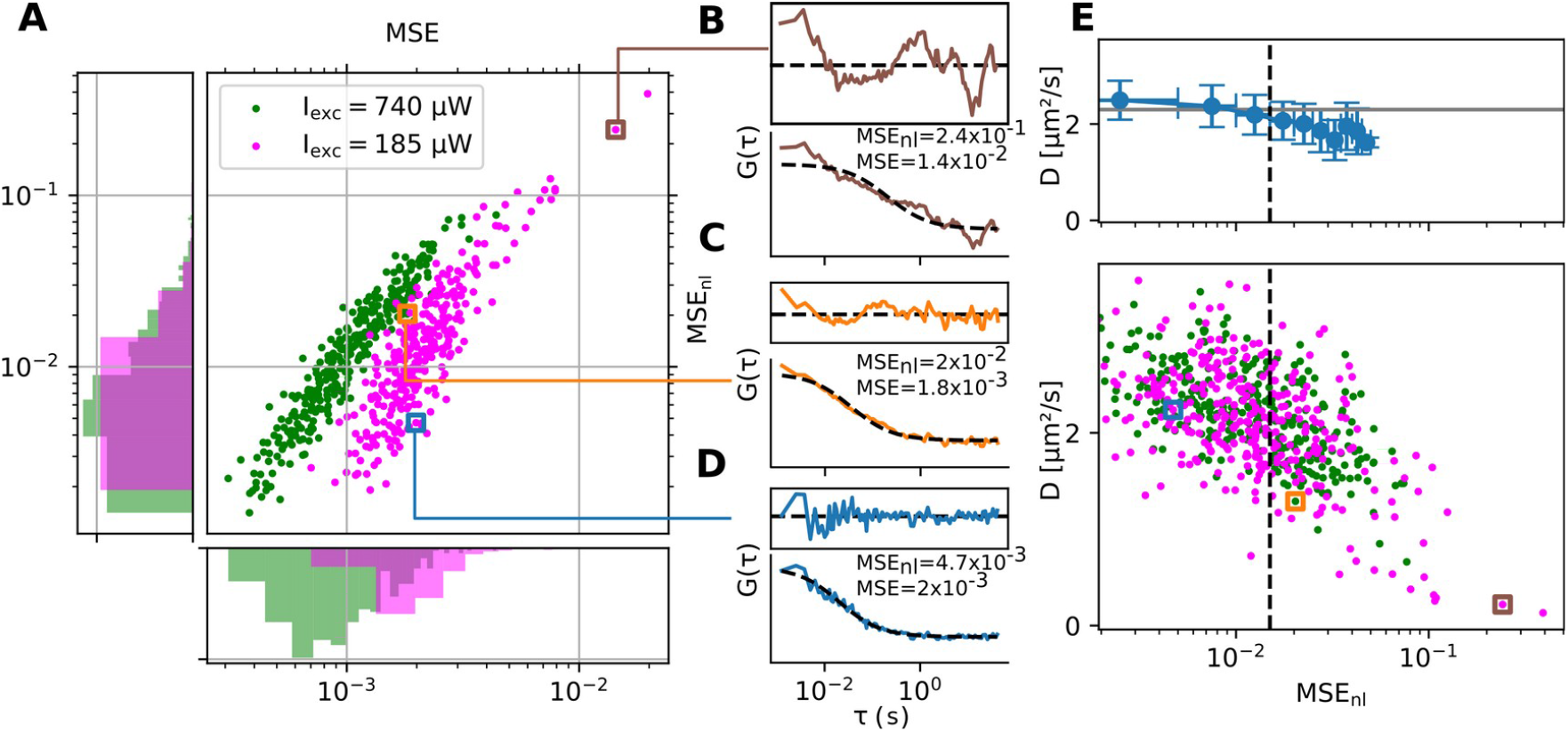
Estimating the fit quality of FCS curves acquired on a DOPC SLB labelled with PE-Rhod using MSE and MSE_nl_. (A) Scatter plot (center) of MSE and MSE_nl_ metrics for each curve acquired in TIR-FCS at low (magenta) and high (green) excitation powers as indicated in the legend, and histograms of the corresponding MSE (bottom) and MSE_nl_ (left) distributions. Colored squares refer to FCS curves shown in panels (B-D). (B-D) Normalised FCS curves (color) and fits (dotted black lines) with (top) fitting residuals, with high MSE and MSE_nl_ (B), medium MSE and high MSE_nl_ (C) and medium MSE and low MSE_nl_ (D). The same scale was used to plot all fitting residuals. (E) Bottom: Scatterplot of measured diffusion coefficient with MSE_nl_ and empirical threshold on fitting quality (dotted vertical line) to discard artefactual FCS curves. Squares represent FCS curves shown in (B-D). Top: Mean +/-std of diffusion coefficient within the MSE_nl_ range represented by lateral errorbars (blue) and value reported for the diffusion coefficient of PE-Rhod in a DOPC SLB in ref (55).

### Impact of membrane curvature

Three assumptions made when fitting FCS curves with the model in equation 1 were not verified when doing TIR-FCS in bacteria. First of all, eq. 1 assumes that the diffusion within the observation area is occuring on a 2-dimensional flat surface. Indeed, TIR-FCS measures an average transit time in the observation area and then calculates a diffusion coefficient as a ratio between the size of the observation area and the transit time. In eq. 1, it is assumed that the size of the observation area is identical to the size of the area in which molecules diffused (the diffusion area), which is true when imaging a flat surface but is not when imaging a curved surface. In the latter case, the diffusion area is larger than the observation area. Second, it is assumed that intensity fluctuations are only caused by molecules moving across the observation area and Poisson noise. When doing TIR-FCS in a curved membrane, molecules moving laterally also change their axial position which under TIRF excitation determines excitation intensity and therefore induces intensity fluctuations. The third assumption is that the system is open, which means that there is an infinite pool of fluorescent molecules diffusing in an infinite-sized reservoir. This latest assumption is never actually verified but it is a good approximation when the mean square displacement of fluorescent emitters during the time of acquisition is much smaller than the reservoir size. This is not the case for membrane markers in bacteria: the average distance traveled by molecules diffusing at a reasonable 1µm^2^/s speed over the course of 1 min (our usual acquisition time) is 15 µm, larger than the characteristic dimensions of a *B. subtilis* cell (typically 5x1µm for exponentially-growing *B. subtilis* cells in LB at 37°C). To evaluate the potential biases in diffusion coefficient measurements caused by these effects, we simulated diffusion on curved surfaces of finite areas: either on the simplest case of a sphere (that can represent *cocci* e.g *Staphylococcus aureus*) or on a cylindrical vessel like the rod-shaped *B. subtilis*. Diffusion on these 3D surfaces were simulated as a Wiener process. We first generated a uniform distribution of initial positions. Position vectors were updated for each step by adding the cross product of the position vector with a random 3D vector of brownian motion, then normalised (see Supporting Materials and Methods for a complete description of the simulation). From the position of individual emitters determined as trajectories (Figure 2A), we could simulate TIR-FCS experiments having the physical parameters of our setup: framerate (1 ms/frame), PSF size (FWHM of 450 nm), TIRF penetration depth (100 nm). Molecular brightness was set to 20 kHz and simulated diffusion coefficient was set to 1 µm^2^/s. The number of molecules was set to reach a density of 1.6 molecules/µm^2^, with a minimum of 10 molecules per simulation.

**Figure 2:**
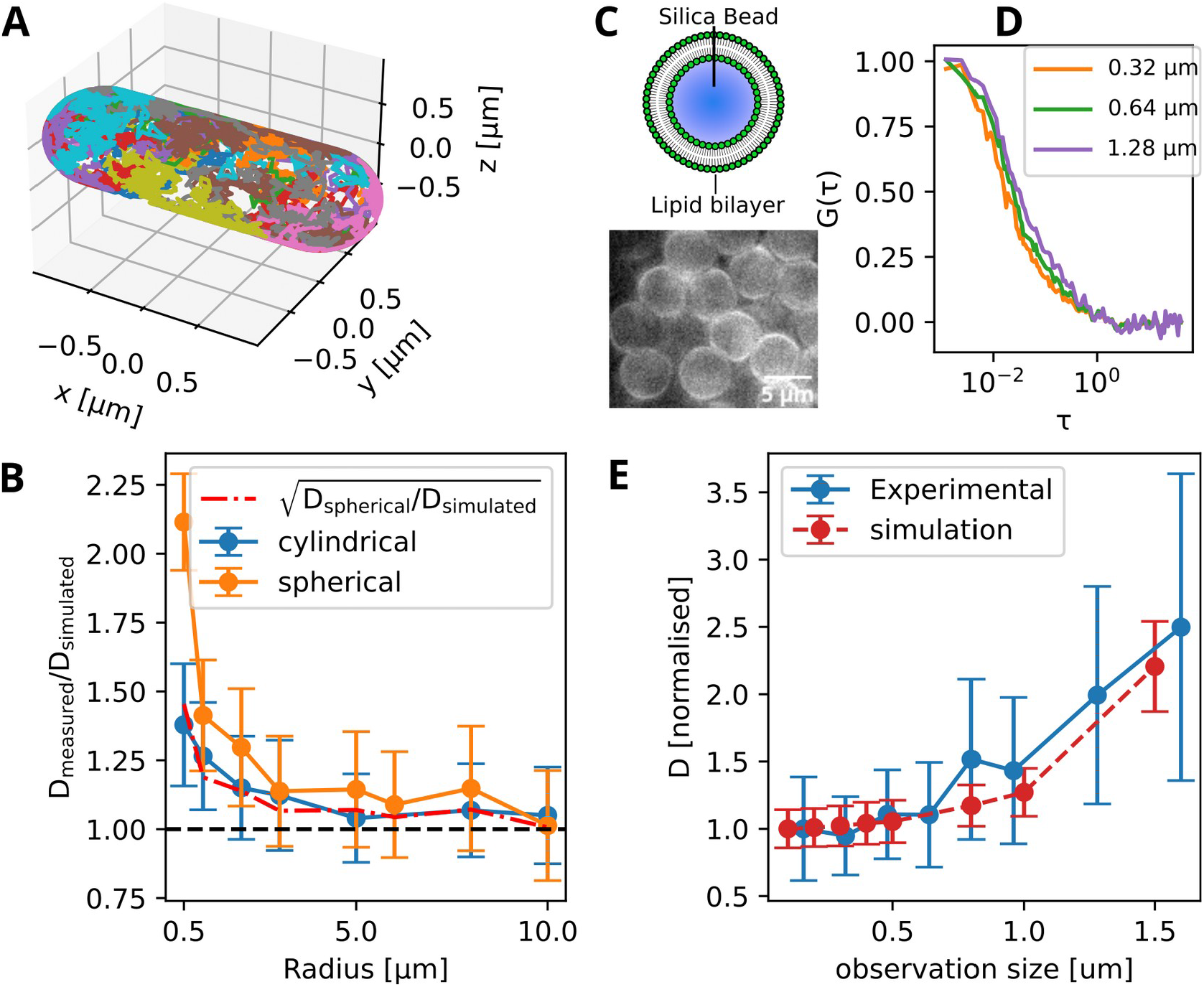
Impact of membrane curvature on TIR-FCS measurements. (A) 3D visualisation of random trajectories generated on a rod (B) Diffusion coefficients measured D_measured_ from simulation of FCS measurements, normalised with simulated value D_simulated,_ in cylindrical (blue) and spherical (orange) coordinates. Dash-dotted red line: square root of normalised diffusion coefficient in a sphere, dotted black line: simulated diffusion coefficient (ground truth). (C) Cartoon of a BSLB (top) and epifluorescence images of BSLBs in an Ibidi chamber (bottom). (D) FCS curves for different observation sizes (obtained by pixel binning) as indicated in the legend, averaged within a 1.28x1.28µm observation area. (E) Diffusion coefficients measured at different observation sizes, normalised with value measured for smallest observation size, from experimental measurements (plain blue line) in 5 µm diameter BSLBs or from simulations in a sphere of same size (dotted red line).

With this simulation framework, we simulated a series of TIR-FCS experiments either on rods (of constant length set to 3 µm) or on spheres of various radii (0.5 to 10 µm) and measured the apparent diffusion coefficient (Figure 2B). We found that in both geometries, the measured diffusion coefficient converged towards the real value for high radius values, which was expected given that a curved membrane of high radius of curvature can be approximated as flat. We found however a significant bias in diffusion coefficient measurement for small radii (below 2 µm). The measurement bias in a rod was well approximated by the square root of the measurement bias in a sphere (Figure 2B, dashdotted line). This can be explained by thinking of curvature as modifying the detection PSF: the detection PSF is modified alongside two dimensions in the case of a sphere and only one dimension in the case of a rod. FCS curves acquired in a rod-shape can therefore be fitted with an updated model based on eq. 1, which is the product of two 1-dimensional fitting models with a fitting bias *f* accounting for curvature alongside one dimension:

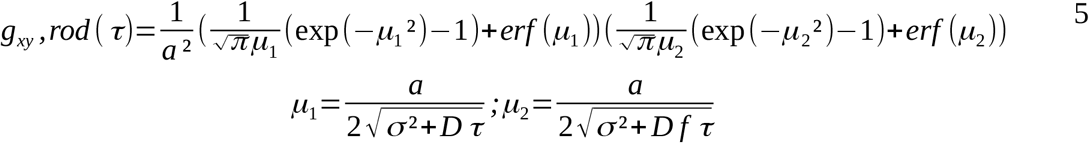

We set the fitting bias *f* to the value of 2.1 for *B. subtilis*, corresponding to the bias induced by a curvature of 500 nm radius on a sphere (Figure 2B). We verified using simulations whether rod length influenced the measurement of diffusion coefficient (Figure S4), and found that for a radius of 500 nm only rod lengths below 2.5 µm affected the diffusion measurements, which is below the lengths of B. subtilis cells we measured in this study (Fig. S5). To validate our simulations experimentally, we used POPC Beads-Supported Lipid Bilayers (BSLB) of 5µm diameter, labeled with the fluorescent lipid PE-Rhod, as a system of 2D diffusion of a controlled diameter (Figure 2C). We performed a series of TIR-FCS experiments in these BSLBs and measured the diffusion coefficient with different observation sizes using different pixel binning values (Figure 2D). The apparent curvature increased as the pixel binning increased and this experiment was therefore a good proxy to measure the effect of different curvatures on measured diffusion coefficient. As expected, we observed an increase in diffusion coefficient with increased observation size, corresponding to an increased effect of curvature on the effective PSF. We observed a similar change in diffusion coefficient with observation size on simulated spheres of identical diameter (Figure 2E). This analysis assumed free diffusion occurring in BSLBs, which was previously shown to not be strictly true (50). Nanoscale hindrances were detected in BSLBs using super-resolution spectroscopy (50), but these were unlikely to affect diffusion measurements at the larger observation scales used here. Indeed, we observed that the measured diffusion coefficient was constant for pixel sizes comprised between 160 and 480 nm (Figure 2E), consistent with a free diffusion approximation.

## Results

### Temperature-induced changes in membrane fluidity

The viscosity of biological membranes heavily depends on the ambient temperature. Reducing temperature increases lipid order, thereby decreasing membrane fluidity, up to the point of phase transition from a fluid membrane to a gel-like structure (56). Poikilothermic (‘cold-blooded’) organisms like bacteria which are naturally exposed to ample changes in temperature adapt their membrane composition in order to maintain fluidity to survive to such changes. Bacterial membrane adaptation to temperature has been widely studied, particularly in the Gram-positive model organism *B. subtilis*, and includes increasing the unsaturated and branched fatty acids. The plasma membrane of steady-state B. subtilis cells contains only low amounts of unsaturated fatty acids (45, 57). Upon cold shock, the enzyme Des desaturates fatty acids (58) immediately (44)(<30 min). Long-term membrane readaptation involves branching instead of unsaturation (45). To measure membrane fluidity of exponentially-growing *B. subtilis* cells in steady-state at 37°C and 20°C, we first used the membrane marker Nile Red, which has the advantage of being bright, photostable, and long-used in *B. subtilis*. Individual diffusion maps alone clearly showed that the diffusion speed of Nile Red was slower at 20°C than at 37°C (Figure 3A-B). This observation was verified through multiple acquisitions, showing an about 2-fold reduction in the diffusion coefficient of Nile red at 20°C relative to 37°C (Figure 3C). We verified that cells were healthy and in exponential phase by monitoring their growth before imaging (Figure S6). Rare cells that were not growing in the microscopy field were excluded from the analysis. We also estimated the impact of phototoxicity by monitoring their growth after imaging and found that cells were growing after imaging, although at a slightly slower rate (Figure S6). To verify whether the diffusion speed of Nile Red was controlled by membrane fluidity and not unforeseen interactions, we performed the same experiment with the membrane dye Di4-ANEPPS (Di4). Diffusion speed of Di4 was slower than Nile Red in similar experimental conditions, which could be expected from its larger size (M=318 g/mol for Nile Red and 480 g/mol for Di4). The reduction of diffusion speed measured between 37°C and 20°C was identical for both dyes (Figure 3C and Table 1), suggesting that it was indeed a change in fluidity that led to the observed reductions in diffusion coefficient. Having established a baseline for the diffusivity of Nile Red in *B. subtilis* at different temperatures, we then sought to observe the remodeling of the membrane in response to a cold shock. For this, we transferred exponentially-growing cells labeled with Nile Red at 37 °C to the microscope at 20°C and performed FCS acquisitions on different cells for 1h. As expected, we observed first a reduction in diffusion coefficient caused by the temperature downshift (Figure 3D-E), then a progressive increase likely caused by membrane adaptation. The ∼30 mins timescale of membrane adaptation we observed was consistent with the significant membrane fatty acids remodeling within 30 mins of cold shock that was previously observed (44). The diffusion coefficients measured were fitted using the model of equation 5 accounting for membrane curvature, assuming a constant radius of 0.5 µm and a cell length larger than 2.5 µm (see Figure S4). We verified that these hypotheses were verified by manual measurements of cell width and length (Figure S5).

**Table 1:** Diffusion coefficients of Nile Red and Di4-ANEPPS measured in B. subtilis at 37°C and 20°C, and ratios of diffusion coefficients at 37°C and 20°C. Presented are the means and standard deviations of the average diffusion coefficients for each acquisition (dots in Figure 3*C*).

| Dye | $D_{37^\circ\text{C}}$ (mean +/- std) | $D_{20^\circ\text{C}}$ (mean +/- std) | $D_{37^\circ\text{C}}/D_{20^\circ\text{C}}$ |
| --- | --- | --- | --- |
| Nile Red | 4.4+/-0.3 | 2.2+/-0.2 | 2+/-0.3 |
| Di4-ANEPPS | 1.9+/-0.1 | 0.9+/-0.07 | 2.1+/-0.3 |

**Figure 3:**
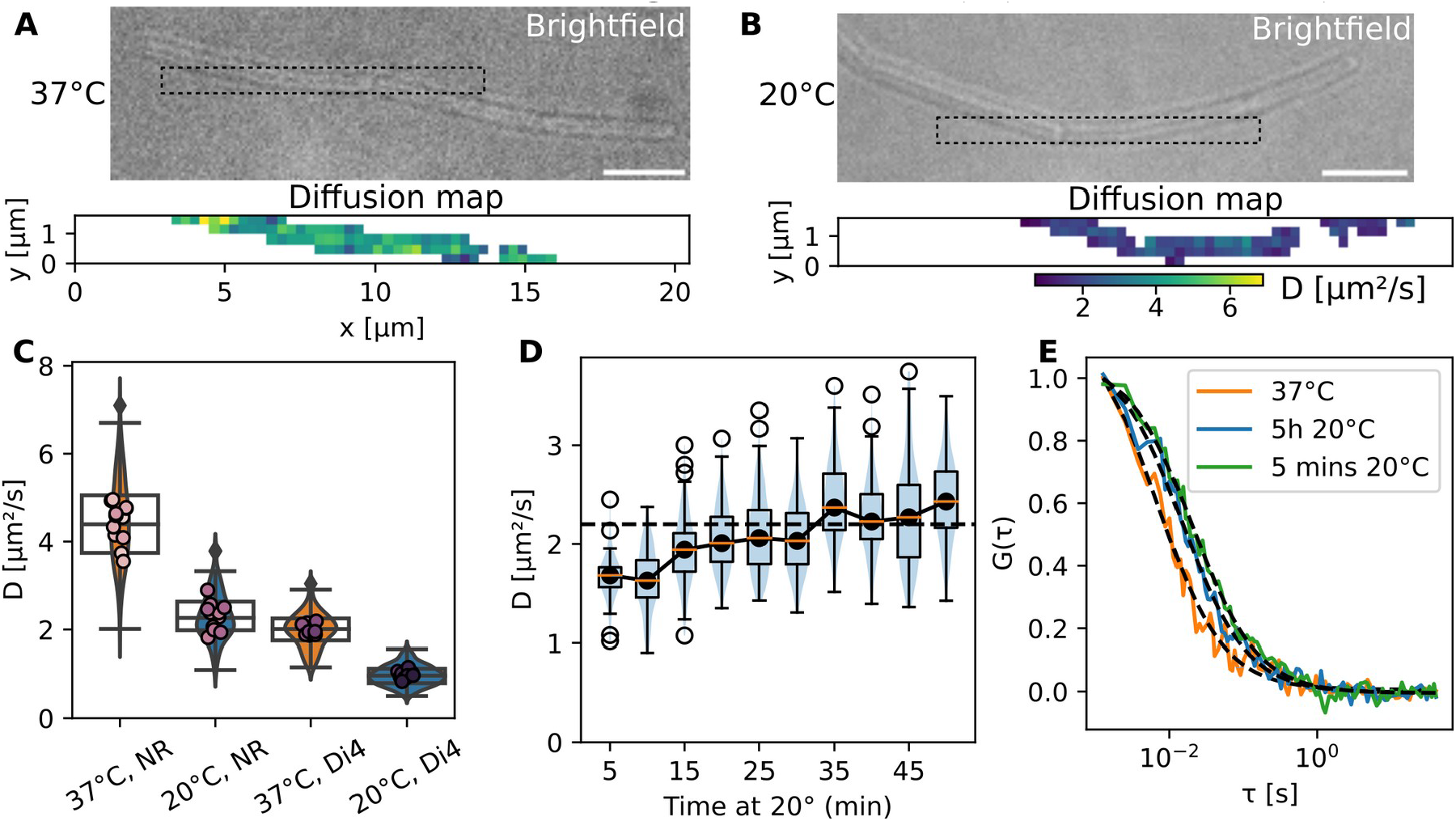
Adaptation of B. subtilis membrane to cold temperature, measured with TIR-FCS. (A-B) representative bright-field (top) with highlighted areas (dotted squares) where TIR-FCS diffusion maps (bottom) of Nile Red in exponentially-growing B. Subtilis were acquired, at 37°C (A) or after 5h at 20°C (B). Scalebars: 5 µm. (C) Diffusion coefficient of Nile Red (left) and Di4 (right) in B. subtilis cells grown and imaged at 37°C (orange) or 5 hours after transfer at 20 °C (blue). Dots: average of individual TIR-FCS acquisitions of one or more cells, 2 biological replicates, n>280 FCS curves per condition. (D) Recovery of diffusion speed of Nile Red upon transfer from 37°C to 20°C, pooling measurements by tranches of 5 mins, n=2 biological replicates. Dotted black line: median diffusion coefficient of Nile Red at 20°C from (C). (E) Normalised FCS curves obtained at 37°C (orange) and 5 mins (green) and 5h (blue) after transfer at 20°C.

## Discussion

In this paper, we demonstrated how TIR-FCS can be used to quantitatively measure membrane fluidity in bacteria, exemplified by the Gram-positive model organism *Bacillus subtilis*. For this, we derived a new fit quality metric that greatly simplified data analysis. Using simulations validated by experiments, we estimated the measurement bias caused by the observation of a curved membrane of finite size in a TIRF field, and used this information to perform unbias measurements of diffusion coefficients in the membrane of *B. subtilis*. We used this assay to measure membrane fluidity, reported by diffusion speed of membrane markers, in the membrane of *B. subtilis* at different temperatures, during membrane remodeling caused by a cold shock and in a steady-state.

The fitting quality metric we derived in the first section is a fine addition to the collection of readily-available quality metrics for FCS quality assessment. Its reliability across datasets with different signal levels might prove useful in future high-throughput FCS studies, whether camera-based or not (59). Using TIR-FCS instead of conventional confocal FCS offered several advantages, some of them already listed: high-throughput measurements, excellent axial selectivity allowing FCS measurements in single membranes and extraction of spatial information. Besides, using an unpolarised evanescent field with TIRF excitation allows excitation of fluorescent molecules irrespective of their dipole orientation. This is a major advantage when working with membrane markers that intercalate within a membrane and keep a fixed orientation, which can be orthogonal to the polarisation plane of e.g confocal excitation and therefore lead to poor excitation quality and inhomogeneous excitation of observation area.

Simulating TIR-FCS in curved surfaces proved very useful for understanding and measuring the potential biases induced by membrane curvature, inhomogeneous excitation of the curved membrane in the evanescent field, and diffusion in a closed system of small size. The respective impact of these three effects was not disentangled by our simulations and further investigations could quantify the individual impact of each of these three effects on the final measurement. Simple 2-dimensional simulations in finite-sized boxes could readily reveal that the small diffusion areas typical of bacterial membranes can lead to substantial biases in measured diffusion speed (Figure S3). Care must be taken when accounting for membrane curvature with simulations to accurately simulate not only the observed geometry, but also the microscope itself as parameters like the PSF size affect the final bias in diffusion coefficient (Figure S4). Furthermore, we did not account for photobleaching in our simulations. This might affect the measurement bias caused by the closedness of the system: in a closed system, the same molecule is more likely to diffuse repeatdly through an observation area than in an open system. However, if there is significant bleaching, a molecule will likely diffuse only once through the observation area and then bleach. We do not expect this effect to significantly change the predictions of our simulations: bleaching was not simulated but occurred in BSLBs, and both systems exhibited a similar behaviour (Figure 2E).

In this work, we showed that the diffusivity of two different membrane markers decreased similarly in response to a decrease in temperature, suggesting a common determinant to their diffusivity: membrane fluidity. Claiming that this assay measures membrane fluidity however requires a couple of reasonable approximations. Membrane fluidity in the field of microbiology is usually referred to as a single parameter (the physical parameter governing diffusion speed in the plasma membrane). However, it is known that the fluidity in the physical sense affects the diffusivity of objects depending on their hydrophobic radius (60). The diffusivity of the membrane markers Nile Red and Di4 is therefore a good proxy for the measurement of membrane fluidity experienced by molecules of comparable size (e.g lipids) but the diffusivity of larger molecules such as transmembrane proteins will be affected by membrane fluidity differently. Relative differences should nevertheless remain the same: the slower diffusion speed of Nile Red at 20°C relative to 37°C indicates that membrane proteins undergoing Brownian motion also diffuse more slowly at 20°C than at 37°C. Furthermore, we did not consider lateral membrane heterogeneities (named functional microdomains in bacteria (61), conceptually equivalent to lipid rafts (62) in eukaryotes) that are thought to recruit certain proteins and lipids in areas of higher molecular order and therefore lower mobility. The diffusivity that we measured in this study likely represents an average of the diffusivity in all domains of the membrane.

Our measurements were performed in the rod-shaped Gram-positive model bacterium B. subtilis. Together with our setup, this offers several advantages: first, the well conserved cell diameter of *B. subtilis* across the cell cycle and growth conditions (63) facilitates correction for membrane curvature. Second, Gram-positive bacteria lack an outer membrane. More care would need to be taken for labeling Gram-negative bacteria to ensure that the used fluorescent marker labels only the membrane of interest, either the inner or the outer membrane. Third, additional care must be taken when performing FCS experiments in Gram-negative bacteria, as the topography of the outer membrane was previously shown to be affected by various compounds (26, 64) which itself affects measurement of diffusion speeds with FCS (65). Our work confirmed the results of previous experiments on the effect of a cold shock on membrane fluidity. Upon cold shock, membrane fluidity decreases, and quickly increases again thanks to a modification of plasma membrane composition that increases unsaturated and branched-chain fatty acids in the membrane (45). It remained unknown by how much membrane fluidity recovered. Fatty acid profiles only provided qualitative information and conflicting results were obtained with environment-sensitive probes (66). DPH anisotropy suggested an incomplete recovery of *B. subtilis* membrane fluidity while fluorescence lifetime measurements of the same probe indicated an identical fluidity at 20 and 37°C (66). Our results allow to settle this debate unambiguously: membrane fluidity does not completely recover after a cold shock in *B. subtilis*: it recovers to the steady-state fluidity at the temperature to which cells were transferred.

Our TIR-FCS assay opens up exciting perspectives in the field of microbiology. The unprecedented ability to quantify membrane fluidity will shed a new light on the biophysics of bacterial membranes and might help to understand severeal cellular processes as well as the mode of action of membrane-targetting antibiotics or antimicrobial peptides. The newly acquired capability to quantify membrane fluidity might also lead to a better understanding of the fundamental role of membrane fluidity in bacterial physiology.

## Author Contributions

A.B., C.B., E.S. and R.C-L. designed the project and wrote the manuscript. A.B. performed the experiment, analysed data and wrote the code.

## Acknowledgements

This project has received funding from the European Research Council (ERC) under the Horizon 2020 research and innovation program (grant agreement No 772178 to R.C.-L.) and under the Marie Sklodowska-Curie grant agreement No 101030628.

## Data availability

Research data will be deposited on Zenodo after publication. Code and manual of the FCS analysis software developed for this project and used throughout this paper can be found at https://github.com/aurelien-barbotin/pyimfcs. Code for simulations can be found at https://github.com/aurelien-barbotin/geomdsim

## Supporting material

**Supp.Figure 1:**
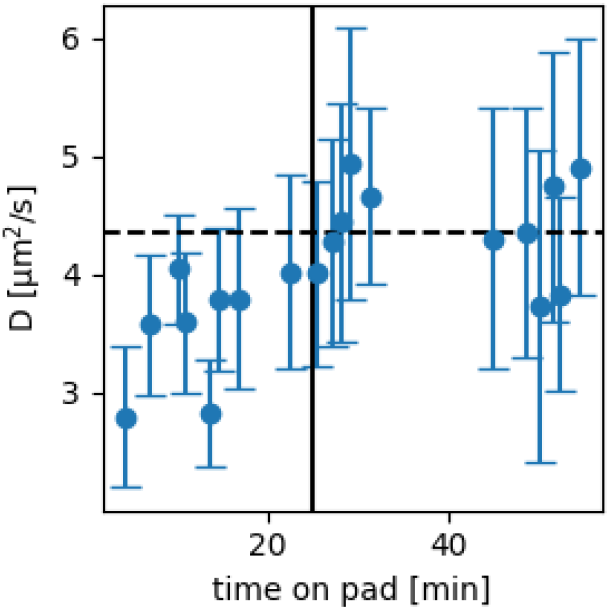
Changes in diffusion coefficient with time spent on pad. Steady-state fluidity is reached after approximately 25 mins (vertical black line), after which imaging can start.

### Simulations

**On a sphere**: First a set of points representing individual fluorescent emitters were distributed randomly on a sphere, as described in (5). The position of each point was described in spherical coordinates. Its azimuthal (θ) and polar (φ) angles were randomly generated using the following equation:

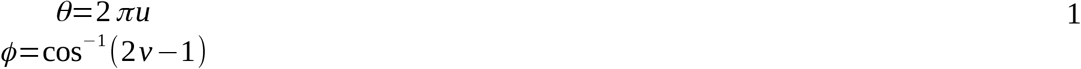

where u and v are drawn from uniform random variables with bounds [0,1[. Trajectories were then converted to cartesian coordinates. At a given time t, the vector position 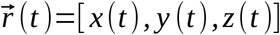 of a point was then updated as follows, as discussed in reference (6):

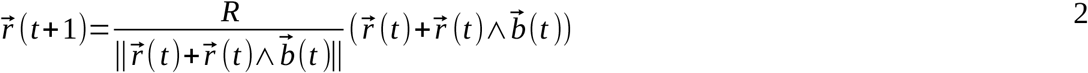

where R is the radius of the sphere and 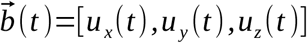 is a three-dimensional random vector drawn from a normal distribution, with each component having a standard deviation of 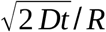, with D the diffusion coefficient. The normalisation factor 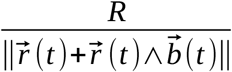 is necessary to keep the vector 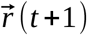 on the surface, as the vector 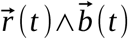 is tangential to the curved surface and therefore 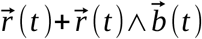 is not on the surface (Figure S2A). Under the conditions that the angle between 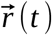 and 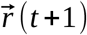 is small, 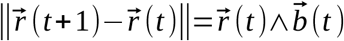 and the simulation of brownian motion is accurate.

**Supp.Figure 2:**
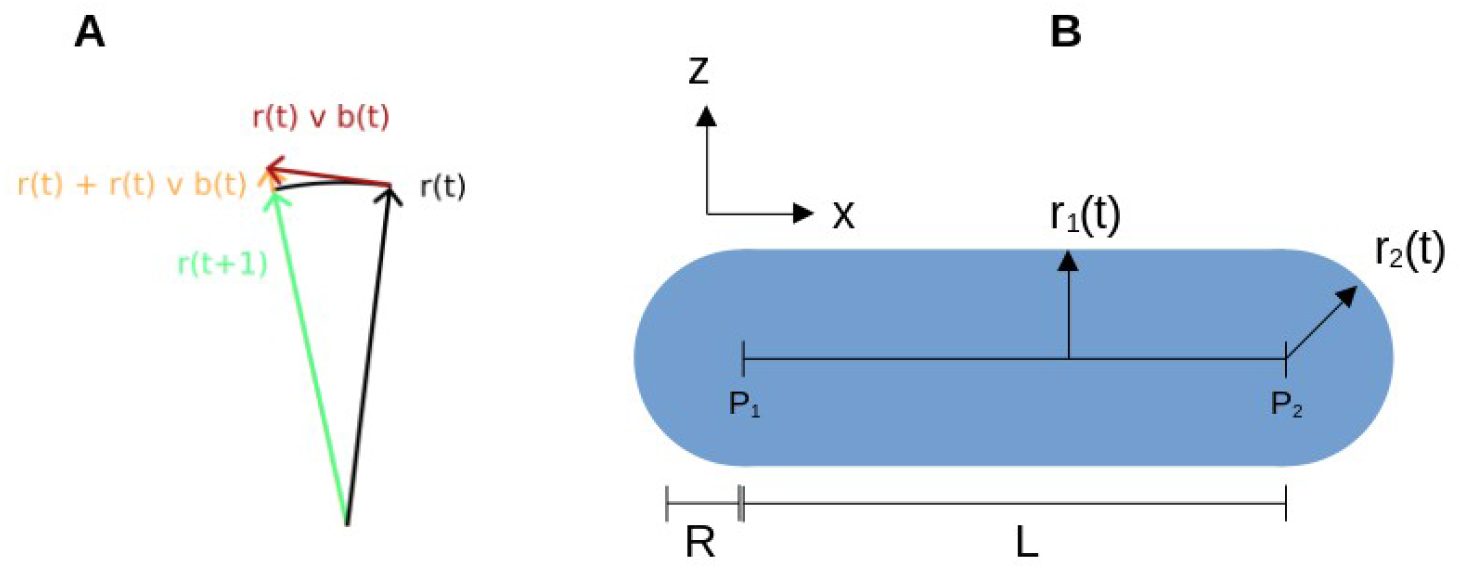
Sketch of the simulation of a Wiener process on curved surfaces. (A): iteration of the Wiener process. B: Sketch of rod-shape simulation.

**On a rod**: the simulation process is very similar. The rod is represented as a cylinder of length L and radius R with two half-spherical parts or radius R at its end (Figure S2B), oriented along the x-axis. The initial distribution of points is done in two steps: a fraction of the total number of the points is distributed on a sphere, while the rest of the points are distributed on a cylinder. The relative fraction of points on the sphere and the cylinder is determined from the relative areas of the spherical and cylindrical parts of the rod. Points drawn on the sphere with a negative x coordinate are moved along the x axis by a distance -L/2, the others by a distance +L/2. The vector position 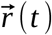 of each point is then iteratively updated following equation 2 as in the previous section, except that in this case the vector 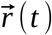 describes the distance to the medial axis (segment [P_1_P_2_] in Figure S2B) of the rod and not to the centre of the sphere. Figure S2B illustrates the two different configurations 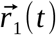 and 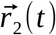 for the vector 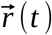.

Parameters used in the simulations of Figure 2 are listed in the following table:

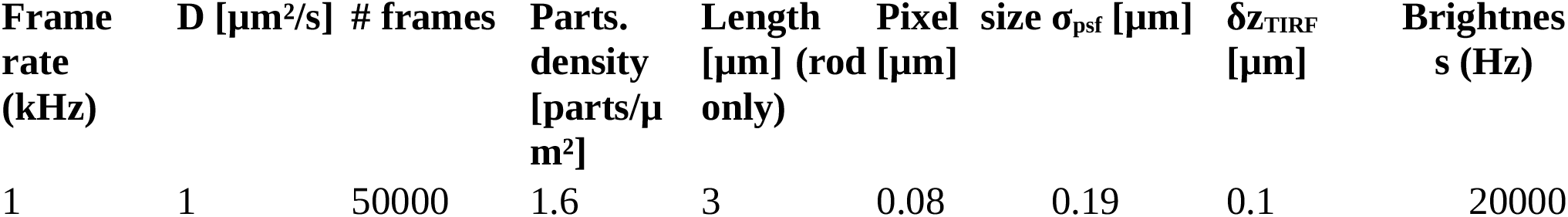

| Frame rate (kHz) | D [$\mu\text{m}^2/\text{s}$] | # frames | Parts. density [parts/$\mu\text{m}^2$] | Length [$\mu\text{m}$] (rod only) | Pixel size [$\mu\text{m}$] | size $\sigma_{\text{psf}}$ [$\mu\text{m}$] | $\delta z_{\text{TIRF}}$ [$\mu\text{m}$] | Brightness (Hz) |
| --- | --- | --- | --- | --- | --- | --- | --- | --- |
| 1 | 1 | 50000 | 1.6 | 3 | 0.08 | 0.19 | 0.1 | 20000 |

Particle density was set to the constant value of 1.6 particle/µm^2^, except in smallest simulations for which it was increased to contain at least 10 particles. TIRF penetration depth δz was defined as:

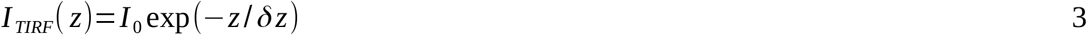

Where I(z) is the depth-dependent TIRF excitation field. To speed up calculations, all particles above 4δz were considered to have a brightness equal to zero and were discarded from the analysis. Analysis was performed with 4x4 binning to an observation area of 320 nm, similar to the one we used in our experiments. An intensity threshold set to 80% of the maximum intensity was also used to analyse simulations as we used in real experiments. Each simulation was performed 9 times. The lateral position of the simulated bacterium was different for each of the 9 simulations to avoid a potential bias.

### Influence of size of closedness of the system

In order to understand if the closedness of the simulated systems described above and in Figure 2 could lead to a bias in diffusion coefficient estimation with imFCS, we simulated a simple system of 2-dimensional Brownian motion in a homogeneous illumination field. Molecules leaving the system on one edge were reintroduced at the corresponding position on the opposite edge (Figure S3A). When the box became very small, we could observe that FCS curves shifted towards shorter lag times and became distorted (Figure S3B). Fitting curves for different box sizes to extract diffusion coefficients confirmed that smaller box sizes, of areas in the order of magnitude of bacterial membrane areas, indeed induced a bias in diffusion coefficient estimation (Figure S3C). It is therefore very likely that part of the measurement biases in Figure 2 were caused by an effect of the small size of the systems observed.

**Supp.Figure 3:**
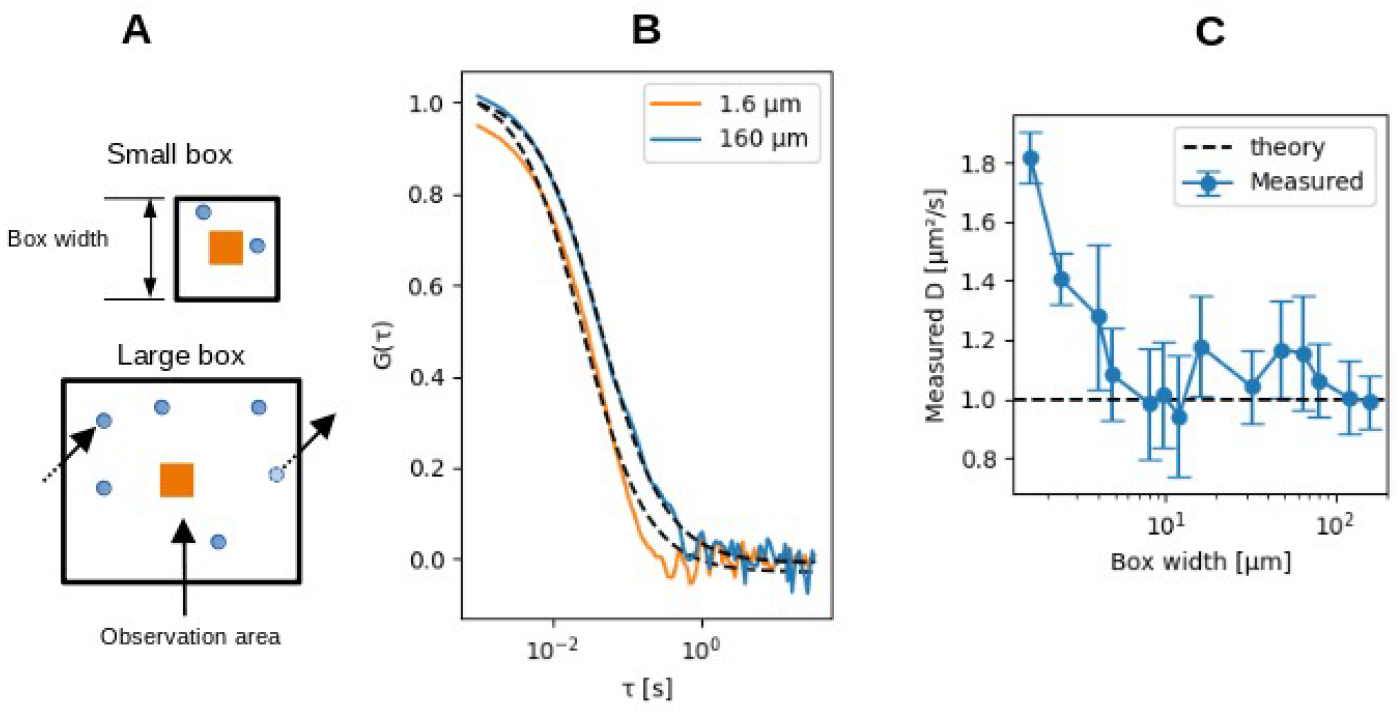
TIR-FCS simulations in a closed box of varying size. (A) sketch of the simulated system with box (black) containing moving particles (blue) and a smaller observation area (centre, orange square) where TIR-FCS is simulated. Top: small box, bottom: large box. A molecule leaving on one side and reentering on the other side is shown with dashed arrows. (B) representative FCS curves obtained when a small (1.6 µm width, orange) and large (160 µm width, blue) simulation box are used. (C) Measured diffusion coefficients in simulated TIR-FCS experiment as a function of box size.

### Influence of rod length and PSF size

**Supp.Figure 4:**
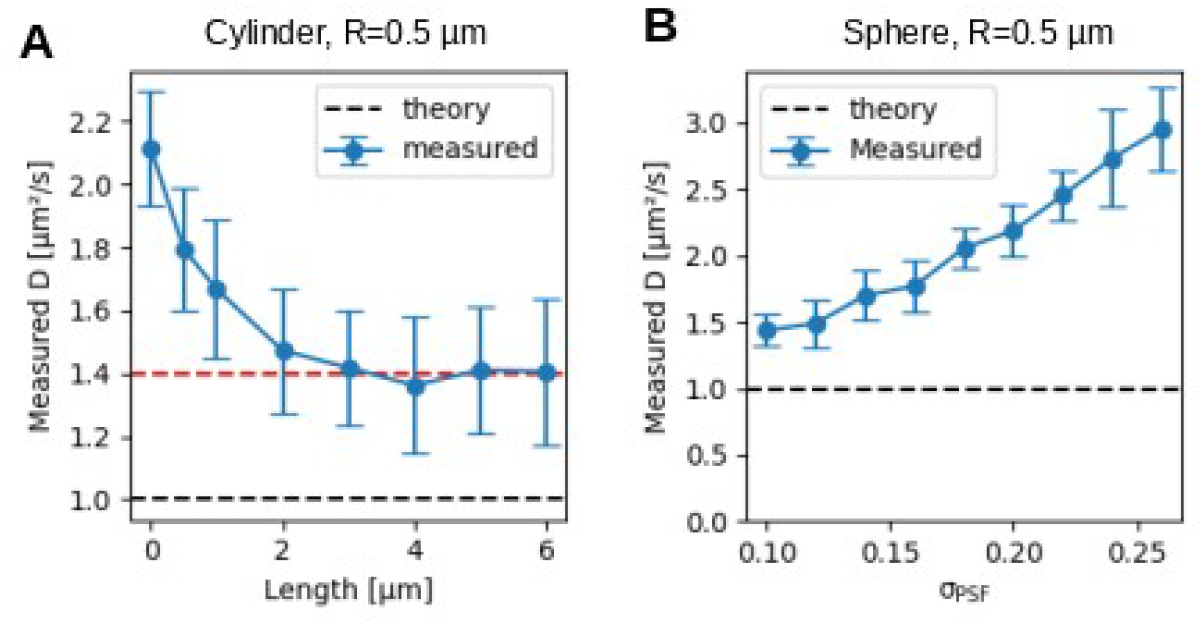
Simulation of the influence of physical parameters on measured diffusion coefficient. (A) Measured diffusion coefficient with length of rod-shape, for a radius of 0.5 µm. Red dashed line: bias for lengths>2.5 µm. (B) Influence of PSF size (parameter σ in equations 1-2) on measured diffusion coefficient, on a sphere of radius 0.5 µm.

## Live bacteria

### Cell morphology in different conditions

Having found that the width and length of bacterial cells bias diffusion coefficient measurements, we verified whether the morphology of *B. subtilis* cells changed between the different experimental conditions investigated here. For this, we acquired epifluorescence images for each of these conditions (Figure S5A-B) and measured cell length and width manually using ImageJ. Cell length was determined by drawing a line between the two poles of each cell and measuring the length of this line. We found that the average cell length was identical at 37°C and immediately after cold shock, decreased after 5hours at 20°C (Figure S5C), but not to a point where cell length biased FCS measurements.

Cell width was measured by plotting the intensity profile alongside a line orthogonal to the cell long axis and measuring the peak-to-peak distance. This method led to an underestimation of the real cell width due to off-axis fluorescence emitted by the top and bottom part of the membrane, hence the relative difference with the well-known diameter of *B. subtilis* of 0.9-1µm (3-4). It revealed however that as expected cell width did not change significantly between experimental conditions and thus that we could apply the same correction factor accounting for membrane curvature to measurements (Figure S5D). Cell width remained constant during cold shock (Figure S5F) and cell length remained well above the 2.5 µm threshold leading to bias in diffusion coefficient (Figure S4A). Panels C and D of Figure S5 were generated using supplementary ref 1.

**Supp.Figure 5:**
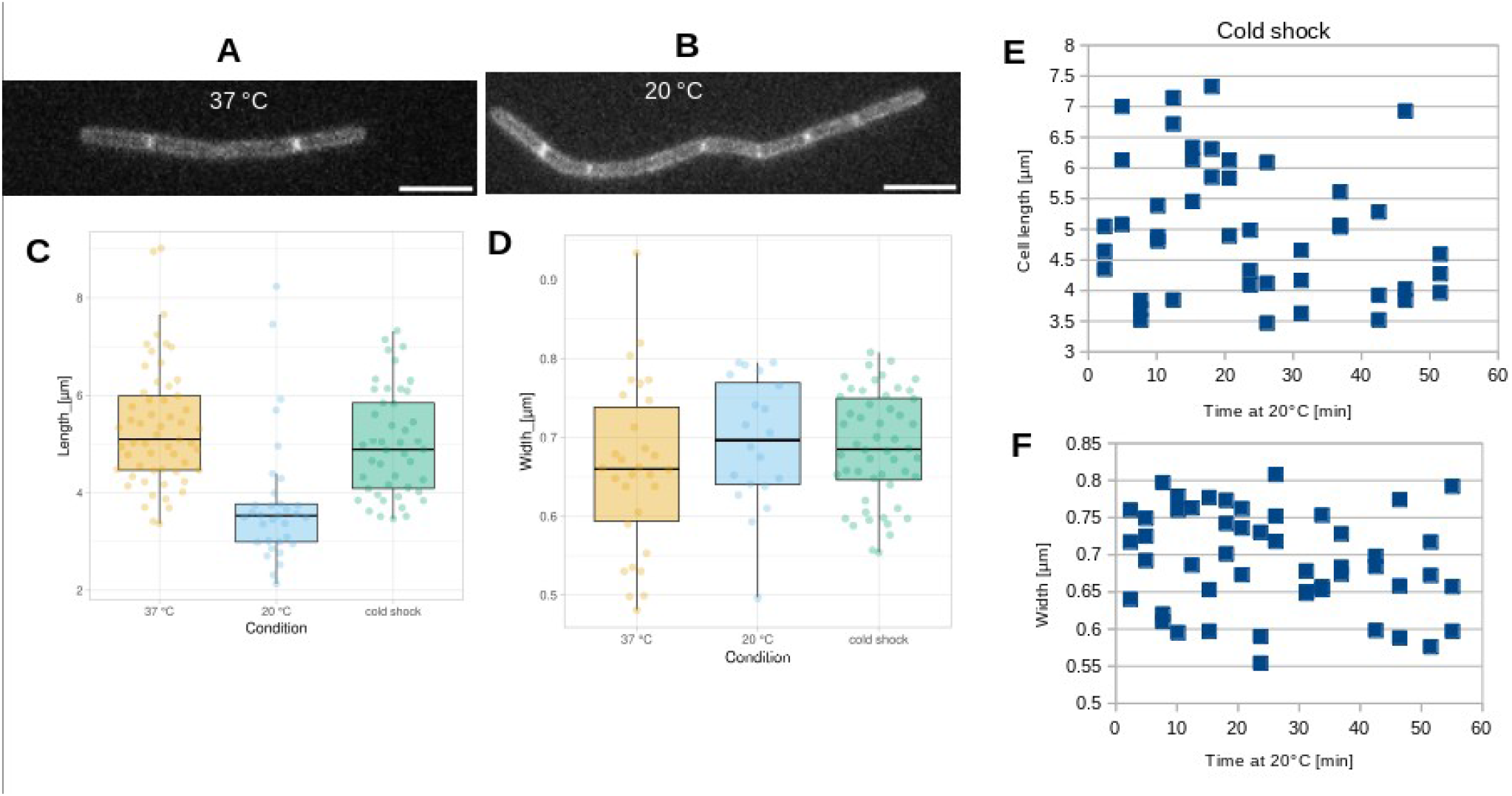
Morphology of B. subtilis at different temperatures. (A-B) epifluorescence images of B. subtilis in exponential phase, labeled with Nile Red, at 37°C (A) and 20 °C (B). Scalebars: 5 µm. (C-D) Length (C) and width (D) of B. subtilis measured from epifluorescence images, in exponential phase at 37°C, 20°C, or during cold shock. (E-F) Scatterplots of cell length (E) and width (F) with time at 20°C immediately after cold shock.

### Impact of FCS measurement on doubling time

Using bright-field timelapses, we verified both cell fitness and the impact of phototoxicity on cell growth. For this, we measured the growth rate of cells used in Figure 3C, at 37°C. We acquired for each chain of cells 3 bright-field images (Figure S6A), one at least 3 mins before the beginning of FCS acquisition, one immediately after the FCS acquisition and one at least 3 mins after FCS acquisition. We measured the length of the cell chain in each bright-field image and calculated doubling times between pairs of frames following the equation (under the assumption of constant cell width as is the case in *B. subtilis*)

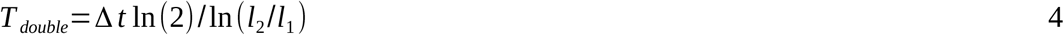

Where *T*_*double*_ is the doubling time, Δ *t* is the time between frames 1 and 2, *l*_1_ and *l*_2_ are the lengths of the cell chain in frames 1 and 2. Cells which doubling was more than twice higher than the nominal doubling time (∼20 mins) were considered not exponentially-growing and therefore excluded from the analysis. Comparing pairwise doubling times before and after FCS (Figure S6B), we found that cells kept growing after FCS, yet at a slightly slower rate, suggesting low photoxicity effects.

**Supp.Figure 6:**
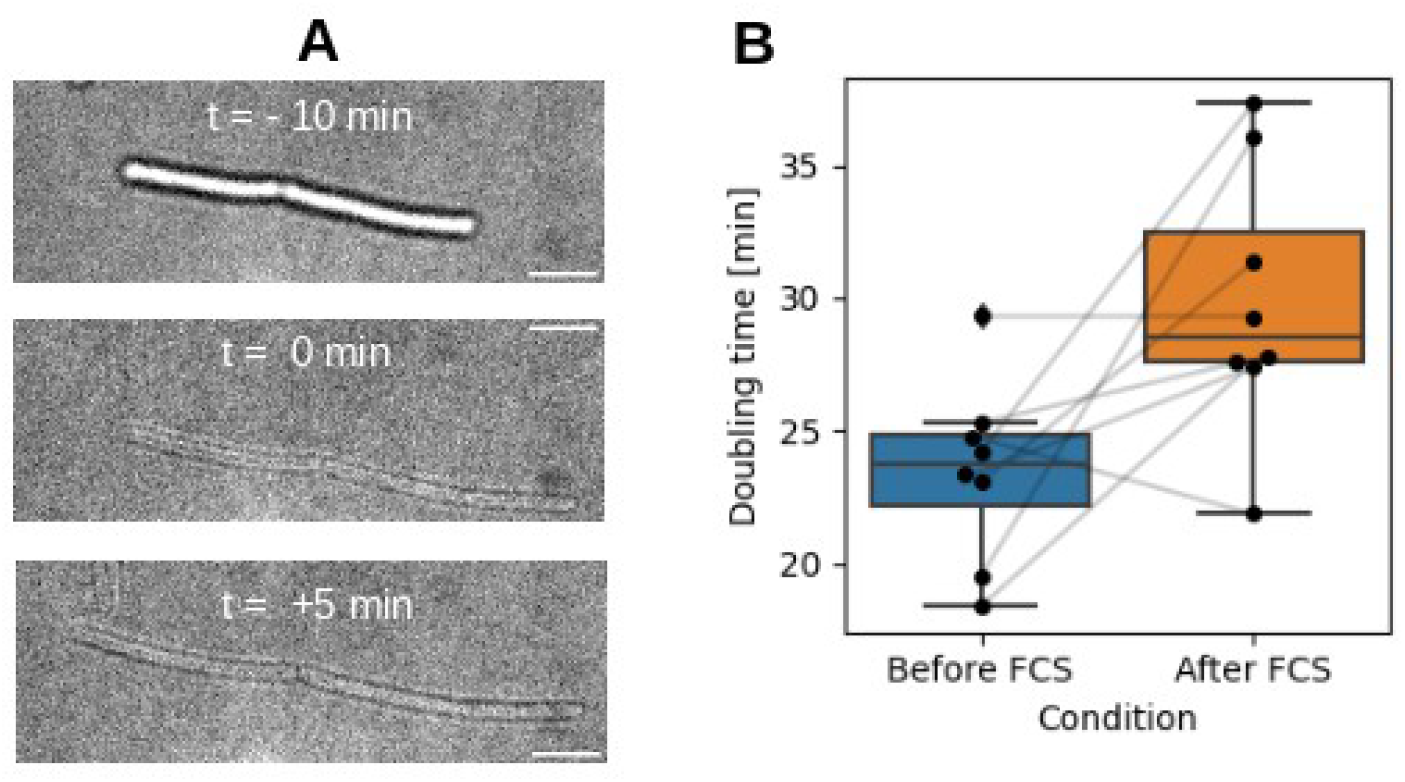
Impact of FCS measurements on the growth rate of Nile Red-labeled B. subtilis cells. (A): bright-field images of growing cells acquired before (top), immediately after (middle) and after (bottom) FCS acquisition. Scale bars: 5 µm. (B) Doubling times calculated from cell elongation, before and after FCS acquisition. Black dots: single doubling times measurements, gray lines link doubling times of the same cell chain.

## Detailed implementation of FCS data processing

### Determination of intensity threshold

In order to avoid biasing diffusion measurements, we needed to exclude TIR-FCS measurements that were too far from the point of contact between the bacterial cell and the coverslip. An efficient way of doing this consisted in removing pixels with an average intensity below a given threshold, as the excitation of the TIRF field decreases with increased distance to the cell centre. To find an appropriate value for this intensity threshold, we plotted a 2D histogram of intensity (normalised with 98^th^ percentile) and diffusion coefficient in 6 acquisitions of exponentially-growing *B. subtilis* labeled with Nile Red at 20°C. We set the threshold to 0.8 so that there was no correlation between diffusion coefficient and intensity (Figure S7).

**Supp.Figure 7:**
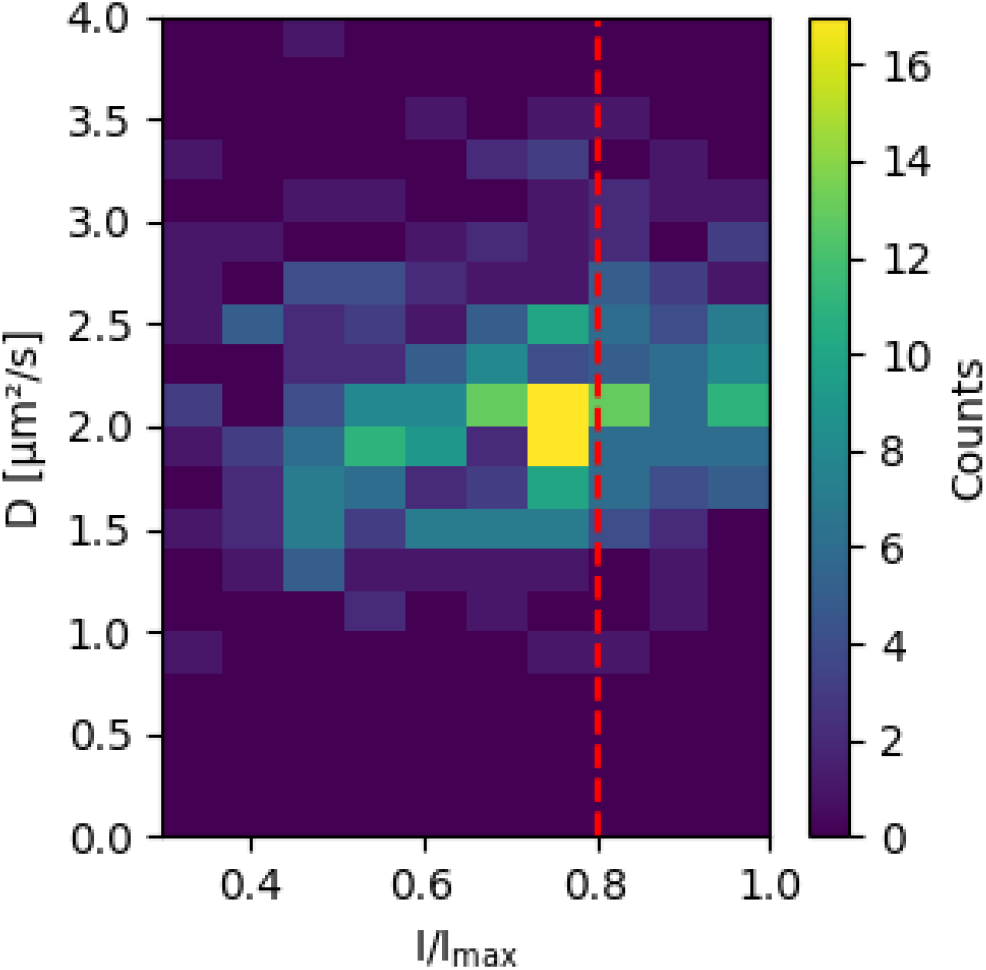
Determination of intensity threshold for unbiased diffusion measurement. Correlation between relative pixel intensity (x axis) and measured diffusion coefficient (y axis) visualised as a 2-dimensional histogram in 6 acquisitions of exponentially-growing B. subtilis at 20°C. Vertical dotted red line: selected threshold

### Bleaching correction and FCS fitting

Bleaching correction was performed using a double exponential fit of the decaying intensity. Intensity timetraces were downsampled 500 times to speed up computations. The resulting traces were fitted with the function:

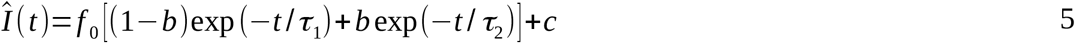

The original intensity timetrace was then corrected as described in ref (2):

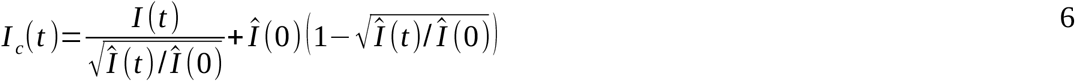

The error function in equation 1 is as :

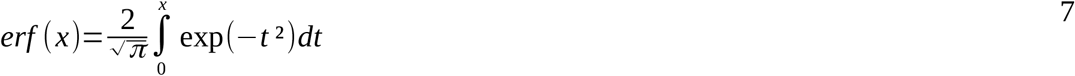

## References

1. Blouin, C.M., Y. Hamon, P. Gonnord, C. Boularan, J. Kagan, C. Viaris de Lesegno, R. Ruez, S. Mailfert, N. Bertaux, D. Loew, C. Wunder, L. Johannes, G. Vogt, F.-X. Contreras, D. Marguet, J.-L. Casanova, C. Galès, H.-T. He, and C. Lamaze. 2016. Glycosylation-Dependent IFN-γR Partitioning in Lipid and Actin Nanodomains Is Critical for JAK Activation. Cell. 166:920–934.

2. Sezgin, E., Y. Azbazdar, X.W. Ng, C. Teh, K. Simons, G. Weidinger, T. Wohland, C. Eggeling, and G. Ozhan. 2017. Binding of canonical Wnt ligands to their receptor complexes occurs in ordered plasma membrane environments. FEBS J. 284:2513–2526.

3. Makarova, M., M. Peter, G. Balogh, A. Glatz, J.I. MacRae, N. Lopez Mora, P. Booth, E. Makeyev, L. Vigh, and S. Oliferenko. 2020. Delineating the Rules for Structural Adaptation of Membrane-Associated Proteins to Evolutionary Changes in Membrane Lipidome. Curr. Biol. 30:367–380.e8.

4. Dewald, A.H., J.C. Hodges, and L. Columbus. 2011. Physical Determinants of β-Barrel Membrane Protein Folding in Lipid Vesicles. Biophys. J. 100:2131–2140.

5. Burgess, N.K., T.P. Dao, A.M. Stanley, and K.G. Fleming. 2008. β-Barrel Proteins That Reside in the Escherichia coli Outer Membrane in Vivo Demonstrate Varied Folding Behavior in Vitro. J. Biol. Chem. 283:26748–26758.

6. Budin, I., T. De Rond, Y. Chen, L.J.G. Chan, C.J. Petzold, and J.D. Keasling. 2018. Viscous control of cellular respiration by membrane lipid composition. Science. 362:1186–1189.

7. Sáenz, J.P., D. Grosser, A.S. Bradley, T.J. Lagny, O. Lavrynenko, M. Broda, and K. Simons. 2015. Hopanoids as functional analogues of cholesterol in bacterial membranes. Proc. Natl. Acad. Sci. 112:11971–11976.

8. Boudjemaa, R., C. Cabriel, F. Dubois-Brissonnet, N. Bourg, G. Dupuis, A. Gruss, S. Lévêque-Fort, R. Briandet, M.-P. Fontaine-Aupart, and K. Steenkeste. 2018. Impact of Bacterial Membrane Fatty Acid Composition on the Failure of Daptomycin To Kill Staphylococcus aureus. Antimicrob. Agents Chemother. 62:e00023–18, /aac/62/7/e00023-18.atom.

9. Vaňousová, K., J. Beranová, R. Fišer, M. Jemioła-Rzemińska, P. Matyska Lišková, L. Cybulski, K. Strzałka, and I. Konopásek. 2018. Membrane fluidization by alcohols inhibits DesK-DesR signalling in Bacillus subtilis. Biochim. Biophys. Acta BBA - Biomembr. 1860:718–727.

10. Popp, P.F., A. Benjdia, H. Strahl, O. Berteau, and T. Mascher. 2020. The Epipeptide YydF Intrinsically Triggers the Cell Envelope Stress Response of Bacillus subtilis and Causes Severe Membrane Perturbations. Front. Microbiol. 11:151.

11. Lee, T.-H., V. Hofferek, F. Separovic, G.E. Reid, and M.-I. Aguilar. 2019. The role of bacterial lipid diversity and membrane properties in modulating antimicrobial peptide activity and drug resistance. Curr. Opin. Chem. Biol. 52:85–92.

12. Loffeld, B., and H. Keweloh. 1996. cis/trans isomerization of unsaturated fatty acids as possible control mechanism of membrane fluidity in Pseudomonas putida P8. Lipids. 31:811–815.

13. Beney, L., and P. Gervais. 2001. Influence of the fluidity of the membrane on the response of microorganisms to environmental stresses. Appl. Microbiol. Biotechnol. 57:34–42.

14. Willdigg, J.R., and J.D. Helmann. 2021. Mini Review: Bacterial Membrane Composition and Its Modulation in Response to Stress. Front. Mol. Biosci. 8:634438.

15. Yoon, Y., H. Lee, S. Lee, S. Kim, and K.-H. Choi. 2015. Membrane fluidity-related adaptive response mechanisms of foodborne bacterial pathogens under environmental stresses. Food Res. Int. 72:25–36.

16. Mendoza, D.D. 2014. Temperature Sensing by Membranes. Annu. Rev. Microbiol. 68:101–116.

17. Zielińska, A., A. Savietto, A. de Sousa Borges, D. Martinez, M. Berbon, J.R. Roelofsen, A.M. Hartman, R. de Boer, I.J. Van der Klei, A.K. Hirsch, B. Habenstein, M. Bramkamp, and D.-J. Scheffers. 2020. Flotillin-mediated membrane fluidity controls peptidoglycan synthesis and MreB movement. eLife. 9:e57179.

18. Nickels, J.D., S. Poudel, S. Chatterjee, A. Farmer, D. Cordner, S.R. Campagna, R.J. Giannone, R.L. Hettich, D.A.A. Myles, R.F. Standaert, J. Katsaras, and J.G. Elkins. 2020. Impact of Fatty-Acid Labeling of Bacillus subtilis Membranes on the Cellular Lipidome and Proteome. Front. Microbiol. 11:914.

19. Strahl, H., F. Bürmann, and L.W. Hamoen. 2014. The actin homologue MreB organizes the bacterial cell membrane. Nat. Commun. 5:3442.

20. Nichols, C.M., J.P. Bowman, and J. Guezennec. 2005. Effects of Incubation Temperature on Growth and Production of Exopolysaccharides by an Antarctic Sea Ice Bacterium Grown in Batch Culture. Appl. Environ. Microbiol. 71:3519–3523.

21. Chattopadhyay, M.K. 2006. Mechanism of bacterial adaptation to low temperature. J. Biosci. 31:157–165.

22. Edgcomb, M.R., S. Sirimanne, B.J. Wilkinson, P. Drouin, and R. Morse. 2000. Electron paramagnetic resonance studies of the membrane fluidity of the foodborne pathogenic psychrotroph Listeria monocytogenes. Biochim. Biophys. Acta. 1463:31–42.

23. Konings, A.W.T., and A.C.C. Ruifrok. 1985. Role of Membrane Lipids and Membrane Fluidity in Thermosensitivity and Thermotolerance of Mammalian Cells. Radiat. Res. 102:86.

24. Saxton, M.J., and K. Jacobson. 1997. SINGLE-PARTICLE TRACKING:Applications to Membrane Dynamics. Annu. Rev. Biophys. Biomol. Struct. 26:373–399.

25. Devkota, R., and M. Pilon. 2018. FRAP: A Powerful Method to Evaluate Membrane Fluidity in Caenorhabditis elegans. Bio-Protoc. 8:e2913.

26. Ponmalar, I.I., J. Swain, and J.K. Basu. 2022. Modification of bacterial cell membrane dynamics and morphology upon exposure to sub inhibitory concentrations of ciprofloxacin. Biochim. Biophys. Acta BBA - Biomembr. 1864:183935.

27. Ragaller, F., L. Andronico, J. Sykora, W. Kulig, T. Rog, Y.B. Urem, D.I. Danylchuk, M. Hof, A. Klymchenko, M. Amaro, I. Vattulainen, and E. Sezgin. 2022. Dissecting the mechanisms of environment sensitivity of smart probes for quantitative assessment of membrane properties. Open Biol. 12:220175.

28. Amaro, M., F. Reina, M. Hof, C. Eggeling, and E. Sezgin. 2017. Laurdan and Di-4-ANEPPDHQ probe different properties of the membrane. J. Phys. Appl. Phys. 50:134004.

29. Poojari, C., N. Wilkosz, R.B. Lira, R. Dimova, P. Jurkiewicz, R. Petka, M. Kepczynski, and T. Róg. 2019. Behavior of the DPH fluorescence probe in membranes perturbed by drugs. Chem. Phys. Lipids. 223:104784.

30. Meacci, G., J. Ries, E. Fischer-Friedrich, N. Kahya, P. Schwille, and K. Kruse. 2006. Mobility of Min-proteins in Escherichia coli measured by fluorescence correlation spectroscopy. Phys. Biol. 3:255–263.

31. Cluzel, P., M. Surette, and S. Leibler. 2000. An Ultrasensitive Bacterial Motor Revealed by Monitoring Signaling Proteins in Single Cells. Science. 287:1652–1655.

32. Dajkovic, A., E. Hinde, C. MacKichan, and R. Carballido-Lopez. 2016. Dynamic Organization of SecA and SecY Secretion Complexes in the B. subtilis Membrane. PLOS ONE. 11:e0157899.

33. Guet, C.C., L. Bruneaux, T.L. Min, D. Siegal-Gaskins, I. Figueroa, T. Emonet, and P. Cluzel. 2008. Minimally invasive determination of mRNA concentration in single living bacteria. Nucleic Acids Res. 36:e73–e73.

34. Diepold, A., E. Sezgin, M. Huseyin, T. Mortimer, C. Eggeling, and J.P. Armitage. 2017. A dynamic and adaptive network of cytosolic interactions governs protein export by the T3SS injectisome. Nat. Commun. 8:15940.

35. Barbotin, A., I. Urbančič, S. Galiani, C. Eggeling, M. Booth, and E. Sezgin. 2020. z-STED Imaging and Spectroscopy to Investigate Nanoscale Membrane Structure and Dynamics. Biophys. J. 118:2448–2457.

36. Kannan, B., J.Y. Har, P. Liu, I. Maruyama, J.L. Ding, and T. Wohland. 2006. Electron Multiplying Charge-Coupled Device Camera Based Fluorescence Correlation Spectroscopy. Anal. Chem. 78:3444–3451.

37. Kannan, B., L. Guo, T. Sudhaharan, S. Ahmed, I. Maruyama, and T. Wohland. 2007. Spatially Resolved Total Internal Reflection Fluorescence Correlation Microscopy Using an Electron Multiplying Charge-Coupled Device Camera. Anal. Chem. 79:4463–4470.

38. Bag, N., D.A. Holowka, and B.A. Baird. 2020. Imaging FCS delineates subtle heterogeneity in plasma membranes of resting mast cells. Mol. Biol. Cell. 31:709–723.

39. Ng, X.W., C. Teh, V. Korzh, and T. Wohland. 2016. The Secreted Signaling Protein Wnt3 Is Associated with Membrane Domains In Vivo: A SPIM-FCS Study. Biophys. J. 111:418–429.

40. Ng, J., R.D. Kamm, T. Wohland, and R.S. Kraut. 2018. Evidence from ITIR-FCS Diffusion Studies that the Amyloid-Beta (Aβ) Peptide Does Not Perturb Plasma Membrane Fluidity in Neuronal Cells. J. Mol. Biol. 430:3439–3453.

41. Bag, N., D.H.X. Yap, and T. Wohland. 2014. Temperature dependence of diffusion in model and live cell membranes characterized by imaging fluorescence correlation spectroscopy. Biochim. Biophys. Acta BBA - Biomembr. 1838:802–813.

42. Yao, Z., and R. Carballido-López. 2014. Fluorescence Imaging for Bacterial Cell Biology: From Localization to Dynamics, From Ensembles to Single Molecules. Annu. Rev. Microbiol. 68:459–476.

43. Wawrezinieck, L., H. Rigneault, D. Marguet, and P.-F. Lenne. 2005. Fluorescence Correlation Spectroscopy Diffusion Laws to Probe the Submicron Cell Membrane Organization. Biophys. J. 89:4029–4042.

44. Konopasek, I., K. Strzalka, and J. Svobodova. 2000. Cold shock in Bacillus subtilis: different effects of benzyl alcohol and ethanol on the membrane organisation and cell adaptation. Biochim. Biophys. Acta. 1464:18–26.

45. Beranova, J., M.C. Mansilla, D. de Mendoza, D. Elhottova, and I. Konopasek. 2010. Differences in Cold Adaptation of Bacillus subtilis under Anaerobic and Aerobic Conditions. J. Bacteriol. 192:4164–4171.

46. Ries, J., S. Chiantia, and P. Schwille. 2009. Accurate Determination of Membrane Dynamics with Line-Scan FCS. Biophys. J. 96:1999–2008.

47. Müller, P. 2012. Python multiple-tau algorithm. .

48. Ries, J., E.P. Petrov, and P. Schwille. 2008. Total Internal Reflection Fluorescence Correlation Spectroscopy: Effects of Lateral Diffusion and Surface-Generated Fluorescence. Biophys. J. 95:390–399.

49. Bag, N., J. Sankaran, A. Paul, R.S. Kraut, and T. Wohland. 2012. Calibration and Limits of Camera-Based Fluorescence Correlation Spectroscopy: A Supported Lipid Bilayer Study. ChemPhysChem. 13:2784–2794.

50. Beckers, D., D. Urbancic, and E. Sezgin. 2020. Impact of Nanoscale Hindrances on the Relationship between Lipid Packing and Diffusion in Model Membranes. J. Phys. Chem. B. 124:1487–1494.

51. Los, D.A., and N. Murata. 2004. Membrane fluidity and its roles in the perception of environmental signals. Biochim. Biophys. Acta BBA - Biomembr. 1666:142–157.

52. Waithe, D., F. Schneider, J. Chojnacki, M.P. Clausen, D. Shrestha, J.B. de la Serna, and C. Eggeling. 2018. Optimized processing and analysis of conventional confocal microscopy generated scanning FCS data. Methods. 140–141:62–73.

53. Ries, J., M. Bayer, G. Csúcs, R. Dirkx, M. Solimena, H. Ewers, and P. Schwille. 2010. Automated suppression of sample-related artifacts in Fluorescence Correlation Spectroscopy. Opt. Express. 18:11073.

54. Schneider, F., P. Hernandez-Varas, B. Christoffer Lagerholm, D. Shrestha, E. Sezgin, M. Julia Roberti, G. Ossato, F. Hecht, C. Eggeling, and I. Urbančič. 2020. High photon count rates improve the quality of super-resolution fluorescence fluctuation spectroscopy. J. Phys. Appl. Phys. 53:164003.

55. Tang, W.H., S.R. Sim, D.Y.K. Aik, A.V.S. Nelanuthala, T. Athilingam, A. Röllin, and T. Wohland. 2023. Deep learning reduces data requirements and allows real-time measurements in Imaging Fluorescence Correlation Spectroscopy. Biophysics.

56. Heimburg, T. 2019. Phase transitions in biological membranes. . pp. 39–61.

57. Suutari, M., and S. Laakso. 1992. Unsaturated and branched chain-fatty acids in temperature adaptation of Bacillus subtilis and Bacillus megaterium. Biochim. Biophys. Acta BBA - Lipids Lipid Metab. 1126:119–124.

58. Mansilla, M.C., and D. de Mendoza. 2005. The Bacillus subtilis desaturase: a model to understand phospholipid modification and temperature sensing. Arch. Microbiol. 183:229–235.

59. Wachsmuth, M., C. Conrad, J. Bulkescher, B. Koch, R. Mahen, M. Isokane, R. Pepperkok, and J. Ellenberg. 2015. High-throughput fluorescence correlation spectroscopy enables analysis of proteome dynamics in living cells. Nat. Biotechnol. 33:384–389.

60. Petrov, E.P., and P. Schwille. 2008. Translational Diffusion in Lipid Membranes beyond the Saffman-Delbrück Approximation. Biophys. J. 94:L41–L43.

61. Bramkamp, M., and D. Lopez. 2015. Exploring the Existence of Lipid Rafts in Bacteria. Microbiol. Mol. Biol. Rev. 79:81–100.

62. Sezgin, E., I. Levental, S. Mayor, and C. Eggeling. 2017. The mystery of membrane organization: composition, regulation and roles of lipid rafts. Nat. Rev. Mol. Cell Biol. 18:361–374.

63. Juillot, D., C. Cornilleau, N. Deboosere, C. Billaudeau, P. Evouna-Mengue, V. Lejard, P. Brodin, R. Carballido-López, and A. Chastanet. 2021. A High-Content Microscopy Screening Identifies New Genes Involved in Cell Width Control in Bacillus subtilis. 6.

64. Manioglu, S., S.M. Modaresi, N. Ritzmann, J. Thoma, S.A. Overall, A. Harms, G. Upert, A. Luther, A.B. Barnes, D. Obrecht, D.J. Müller, and S. Hiller. 2022. Antibiotic polymyxin arranges lipopolysaccharide into crystalline structures to solidify the bacterial membrane. Nat. Commun. 13:6195.

65. Gesper, A., S. Wennmalm, P. Hagemann, S.-G. Eriksson, P. Happel, and I. Parmryd. 2020. Variations in Plasma Membrane Topography Can Explain Heterogenous Diffusion Coefficients Obtained by Fluorescence Correlation Spectroscopy. Front. Cell Dev. Biol. 8:767.

66. Heřman, P., I. Konopásek, J. Plášek, and J. Svobodová. 1994. Time-resolved polarized fluorescence studies of the temperature adaptation in Bacillus subtilis using DPH and TMA-DPH fluorescent probes. Biochim. Biophys. Acta BBA - Biomembr. 1190:1–8.

## Supplementary References

1. Lord, S. J.; Velle, K. B.; Mullins, R. D.; Fritz-Laylin, L. K. SuperPlots: Communicating Reproducibility and Variability in Cell Biology. Journal of Cell Biology 2020, 219 (6), e202001064. 10.1083/jcb.202001064.

2. Ries, J., S. Chiantia, and P. Schwille. 2009. Accurate Determination of Membrane Dynamics with Line-Scan FCS. Biophysical Journal. 96:1999–2008.

3. Errington J, Aart LTV. 2020. Microbe Profile: Bacillus subtilis: model organism for cellular development, and industrial workhorse. Microbiology (Reading). May;166(5):425–427.

4. Juillot, D., C. Cornilleau, N. Deboosere, C. Billaudeau, P. Evouna-Mengue, V. Lejard, P. Brodin, R. Carballido-López, and A. Chastanet. 2021. A High-Content Microscopy Screening Identifies New Genes Involved in Cell Width Control in Bacillus subtilis. 6.

5. https://www.bogotobogo.com/Algorithms/uniform_distribution_sphere.php

6. https://math.stackexchange.com/questions/3725288/infinitesimal-generator-of-the-brownian-motion-on-a-sphere

